# Origin of diverse phosphorylation patterns in the ERBB system

**DOI:** 10.1101/2020.09.25.313965

**Authors:** T. Okada, H. Miyagi, Y. Sako, M. Hiroshima, A. Mochizuki

## Abstract

Intercellular signals induce various cellular responses, including growth, proliferation, and differentiation, via the dynamic processes of signal transduction pathways. For cell-fate decisions, ligand-binding induces the phosphorylation of ERBB receptors, which in turn activate downstream molecules. The ERBB family includes four subtypes, which diverged through two gene duplications from a common ancestor, of different kinetic properties. Differences in the expression patterns of the subtypes have been reported between different organs in the human body. However, how these different kinetic and expression properties influence the diverse phosphorylation dynamics of ERBB proteins is not well understood. Here we study the phosphorylation dynamics by experimental and mathematical analyses. The experimental measurements clarified that the phosphorylation dynamics heavily depend on the ERBB expression profiles. We developed a mathematical model consisting of the four subtypes as monomers, homodimers and heterodimers and estimated the rate constants governing the phosphorylation dynamics from the experimental data. To understand the origin of the diversity of the phosphorylation responses of the ERBB system, we analyzed the expression profiles and reaction rates of the ERBB subtypes. The difference in phosphorylation rates between ERBB subtypes showed a much larger contribution to the diversity of the phosphorylation responses than did the dimerization rates. This result implies that divergent evolution in phosphorylation reactions rather than in dimerization reactions after whole genome duplications was essential for increasing the diversity of the phosphorylation responses.

**Statement of Significance:** It is known that the expression patterns of a protein family are different between different organs in the human body. This difference is considered essential for the functional diversity between organs. However, the dynamical processes that translate expression patterns into function are not well understood. Here we study the origin of the diversity in ERBB phosphorylation patterns by combining experimental and mathematical methods. We confirmed experimentally that phosphorylation patterns depend on the ERBB expression profiles. Our mathematical model found that differences in phosphorylation rates made the largest contribution to the diversity of phosphorylation patterns between ERBB subtypes.

## Introduction

Signal transduction pathways govern the cellular response to an intercellular signal so as to affect cell growth, proliferation, differentiation and other processes. In the ERBB-RAS-MAPK system (1), which is responsible for cell-fate decisions, more than 100 biomolecules are connected via chemical reactions or regulations of the reactions to constitute a complex network (2). The ERBB family includes four subtypes, ERBB1, ERBB2, ERBB3, and ERBB4, which are all transmembrane proteins and receptor tyrosine kinases (3–5). ERBB1 is also called epidermal growth factor receptor (EGFR), coming from the name of its ligand (6).

ERBB proteins have more than 10 types of extracellular ligands (7) that are all small peptides produced by neighboring cells. Each ERBB paralog exhibits overlapping selectivity and specificity to a subset of ligand species that stimulates a given biological function. In general, ligand-binding induces ERBB phosphorylation, which in turn activates the MAPK, PI3K-AKT, and other pathways in parallel (8). In other words, ERBB family proteins act as “gatekeepers” of signal transduction pathways.

Ligand-binding causes ERBB proteins to form homodimers or heterodimers (9, 10). Phosphorylation of the protomers in the ERBB dimers takes place on the tyrosine residues in the cytoplasmic domain (Fig. 1a), triggering the activation of various cytoplasmic proteins in the signaling pathways (signal transduction), and finally altering the cell structure and function.

**Fig. 1.**
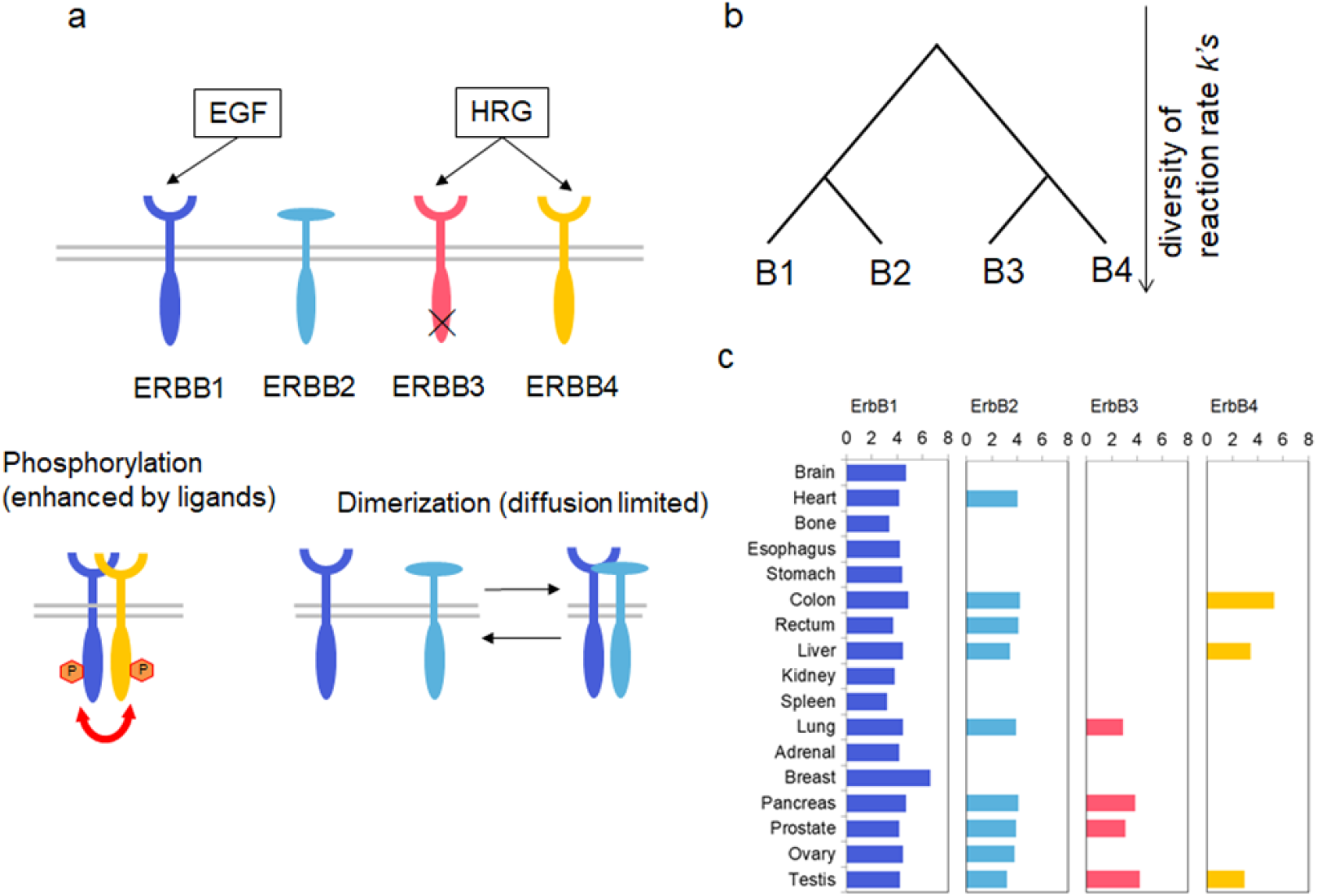
**a.** Schematic of the reactions in the ERBB system. **b**. Schematic phylogenic tree of the ERBB family. **c**. ERBB expression profiles from various human organs. Values are taken from ProteomicsDB (https://www.proteomicsdb.org/).

The phosphorylation level of each ERBB protein depends on the combination of ERBB species in the dimer pair, and differences in the phosphorylation levels influence the downstream signaling. To understand the role of the ERBB network in signaling pathways totally, we need to understand the phosphorylation dynamics as a dynamical system consisting of multiple ERBB subtypes of diverse properties.

In vertebrates other than teleostei, the ERBB family consists of four members, ERBB1-4. These four members are paralogs that diverged with two gene duplications from a common ancestor (Fig. 1b) (11–13). The first divergence produced two members, the ancestor of ERBB1 and ERBB2 and that of ERBB3 and ERBB4. The present four members appeared in the second divergence. After each divergence, the ERBB family members are thought to have acquired unique protein properties, including different rate constants for ligand-binding, dimerization, and phosphorylation.

In addition to the different kinetic properties, different expression levels between ERBB subtypes should influence the phosphorylation dynamics. It has been reported that different organs in the human body show different ERBB expression profiles (Fig. 1c) (14–16). Further, the different expression patterns may be the origin of the diversity of the responses of ERBB signaling pathways between organs.

In this paper, we study the phosphorylation dynamics of ERBB systems consisting of the four subtypes as monomers, homodimers and heterodimers by combining experimental measurements and mathematical analyses. We measured the phosphorylation responses to two ligands from three cell lines and found the phosphorylation dynamics highly depended on the ERBB expression profile. The mathematical modeling was used to estimate and identify differences in the reaction rate constants governing the phosphorylation dynamics between ERBB subtypes. We examined the diversity of the phosphorylation responses; expression profiles were varied in a wide range and the number of different phosphorylation patterns was counted. We found that the diversity of the phosphorylation rates between ERBB subtypes had a much larger contribution to the diversity of the phosphorylation responses than the diversity of the dimerization rates did. This finding implies that divergent evolution in phosphorylation reactions rather than dimerization reactions after whole genome duplications was essential for increasing the diversity of the phosphorylation responses among subtypes.

## Materials and Methods

### Time-lapse measurements

HeLa, A431, and MCF7 cells were cultured in DMEM (WAKO #044-29765) containing 10% FBS to high confluence in 35 mm dishes under 5% CO_2_ at 37 °C. One day before the experiment, the cells were starved in DMEM without FBS and phenol red. The cells were stimulated with 20 nM EGF (PEPROTECH) or 30 nM HRG (R&D Systems) and incubated for 5, 10, 15, 30, and 60 min at room temperature (r.t.). At each time point, the cells were washed with PBS and lysed in 100 uL of 1x SDS sample buffer containing 1 mM Na_3_VO_4_ to avoid dephosphorylation during the sample preparation. The collected samples were boiled at 95 °C for 30 min and then cooled on ice, and SDS-PAGE separation was executed in a 7.5–10% polyacrylamide gel. The separated fractions were transferred to a membrane using a semi-dry blotting apparatus. After blocking the membrane with 2.5% skim milk in Tris buffered saline with 0.1% Tween 20, the primary antibody reaction was carried out at 4 °C overnight, in which anti-phospho-ERBB antibodies for pY1173 of ERBB1 (CST #4407), pY1221/1222 of ERBB2 (CST #2243), pY1289 of ERBB3 (CST #4791), and pY1284 of ERBB4 (CST #4757) were diluted to 1:300–1:500 and applied on the membrane. Then, incubation with a 1:1000 secondary antibody (anti-rabbit IgG) conjugated with horse radish peroxidase (HRP, CST #7074) was done at r.t. for 1 hour. Chemical luminescence from the second antibody reacted with ECL prime reagent (GE Healthcare) was detected by a lumino image analyzer, ImageQuant LAS500 or LAS4000 (GE Healthcare).

### ERBB expression profiles

The relative expression levels of ERBB proteins in the cells were estimated from western blotting. For quantitative comparisons, correction factors compensating for differences in the antibody titers were determined from western blotting ERBBs fused with GFP (ERBB-GFP), i.e., the expression of each type of ERBB-GFP was measured in the same extract of cells with anti-ERBB and anti-GFP (CST #2956) antibodies, and the titer of each anti-ERBB antibody was normalized against that of anti-GFP antibody (Fig. S1). Crosstalk between anti-ERBBs was observed as ERBB1 detected by anti-ERBB4 antibody, however, in most cases, the difference in the molecular weight between ERBBs allowed for the detection of specific staining. In MCF7 cells, the missed staining of ERBB4 (Fig. 3a) might be because of an overlap with the minor upper band of the anti-ERBB1 staining. We excluded the overlapped band in the estimation to minimize the detection error, because the ERBB4 expression was relatively high, thus, the estimated lower-limit value was used in the mathematical analysis described below.

### Overexpression experiments

A gene transfer plasmid that included the sequence of ERBB1, ERBB2, or ERBB3 was transfected to induce the overexpression of the given ERBB protein. The transfection was carried out with a combination of DNA and reagent product that maximized the expression level of transfected ERBB: for HeLa, A431, and MCF7 cells, this combination was 10, 7.5, and 5 μg DNA/35-mm dish and the lipofection reagents of Lipofectamine 3000, Lipofectamine LTX&PLUS, and Lipofectamine 2000 (ThermoFisher), respectively. Two dishes of each transfectant were prepared and cultured for 1 day. One dish was used for western blotting to measure the phosphorylation and expression levels of each ERBB, and the other was used to check the fraction of ERBB-overexpressing cells by fluorescence immunostaining. For the latter assay, cells were cultured on a coverslip in the dish for observation under a fluorescence microscope. In the western blot analysis, only the cultures containing > 90% of ERBB-overexpressing cells were used. To detect the total amount of ERBBs, anti-ERBB antibodies were used with dilutions of 1:500 for ERBB1 (SANTA CRUZ #sc-03), 1:300 for ERBB2 (CST #4290), 1:500 for ERBB3 (SANTA CRUZ #sc-285), and 1:300 for anti-ERBB4 (SANTA CRUZ #sc-283). The secondary antibodies were 1:2000 diluted HRP-linked anti-IgG antibodies (CST #7074 for rabbit and CST #7076 for mouse). ECL prime reagent (GE) was used for acquiring the chemiluminescence signals. For the fluorescence immunostaining, cells without ligand stimulation were fixed by –20 °C MeOH for 2 min, washed with HBSS 3 times and left at 4 °C overnight. After the blocking process with HBSS containing 2% BSA for 15 min, 1:300 diluted anti-ERBB1, anti-ERBB2, or anti-ERBB3 was used for the primary antibody incubation for 1 hour at r.t. Coverslips were washed three times with HBSS and then incubated with 1:1000 anti-IgG antibody conjugated with Alexa 488 (ThermoFisher A-11034) for 1 hour at r.t. The secondary antibody-labeled samples were observed under an inverted fluorescence microscope (Nikon Ti) with a 10X objective lens (Nikon PalnApo) and a filter set suitable for Alexa488. Image acquisition was carried out through a sCMOS camera (ORCA Flash 2.0, HAMAMATSU) with NIS elements software (Nikon).

### Quantification of western blot staining

The intensities of the bands detected in the western blotting were quantified using ImageJ software (NIH). Rectangular ROIs were set in regions of the signal (band) and background; the latter was located far enough from the former so that there was no overlap. The signal intensity was defined as the difference in the average intensities of the two regions.

## Experimental results

### Phosphorylation of ERBB

The phosphorylation of ERBB subtypes (ERBB1, ERBB2, ERBB3, and ERBB4) was detected semi-quantitatively in A431, HeLa, and MCF7 cells stimulated with epidermal growth factor (EGF) or heregulin (HRG) (Fig. 2). These cells are known to show different responses to the same ERBB ligands. We stimulated the cells with EGF, which is a ligand of ERBB1, or HRG, a ligand for ERBB3 and ERBB4, with a saturation concentration of 20 and 30 nM, respectively. At the present time, no intrinsic ligand is known for ERBB2. The ligand concentration was changed in a step-wise manner. The initial response of the ligand-induced phosphorylation was quantified by western blotting at several time points within 60 min of the ligand application.

**Fig. 2.**
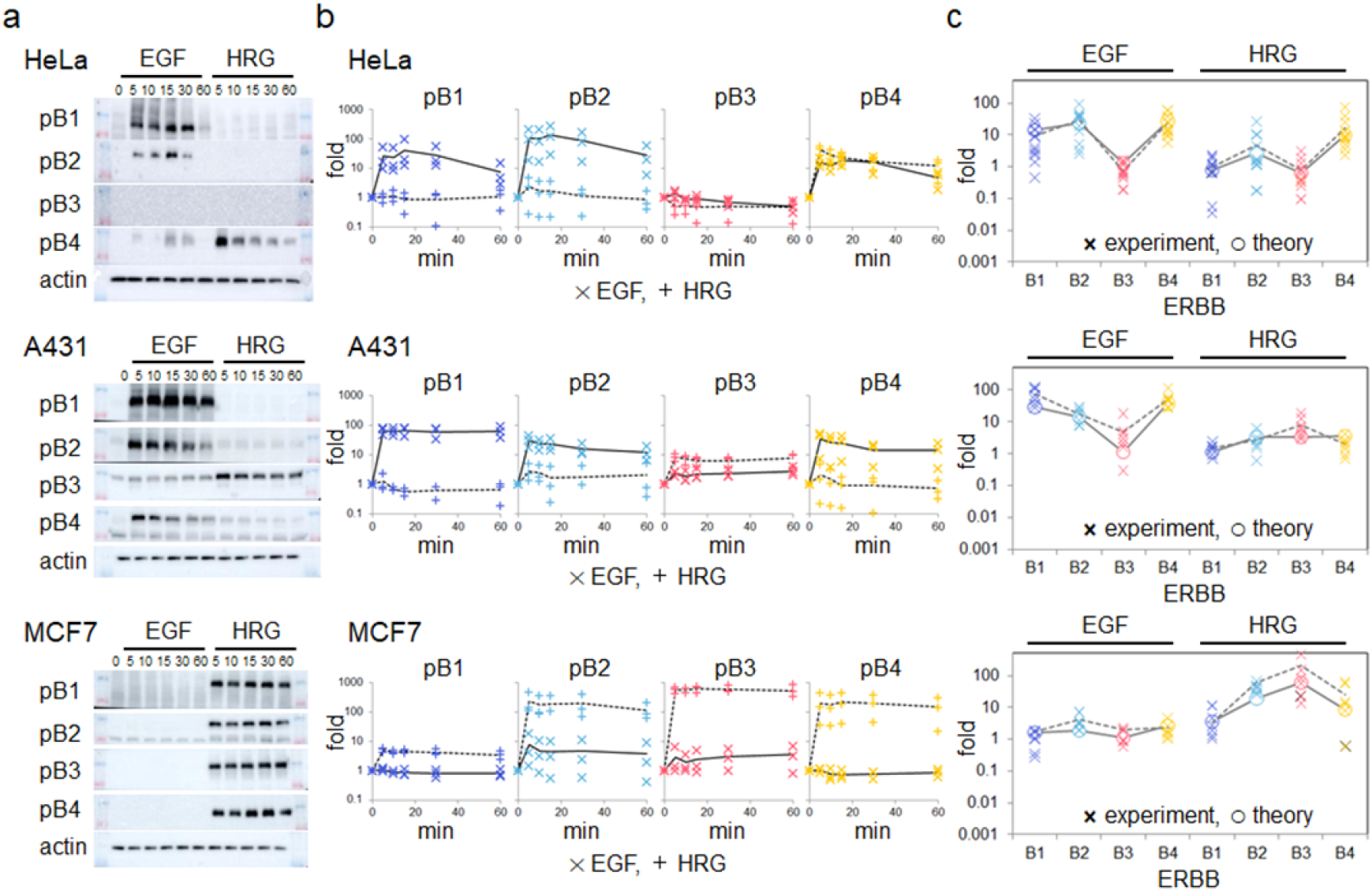
Phosphorylation of ERBB induced by EGF or HRG ligands in HeLa, A431, and MCF7 cells. **a**. Western blotting of ERBB1-B4 after ligand stimulation (numbers indicate minutes). **b**. The time course of the ERBB phosphorylation obtained from the western blotting. Fold changes compare the phosphorylation level to just before the ligand stimulation (time 0). Closed circles and open triangles indicate EGF and HRG stimulation, respectively. **c**. The fold change of ERBB phosphorylation at 5 min. Crosses and open circles respectively represent the experimental results and global model fitting.

**Fig. 3.**
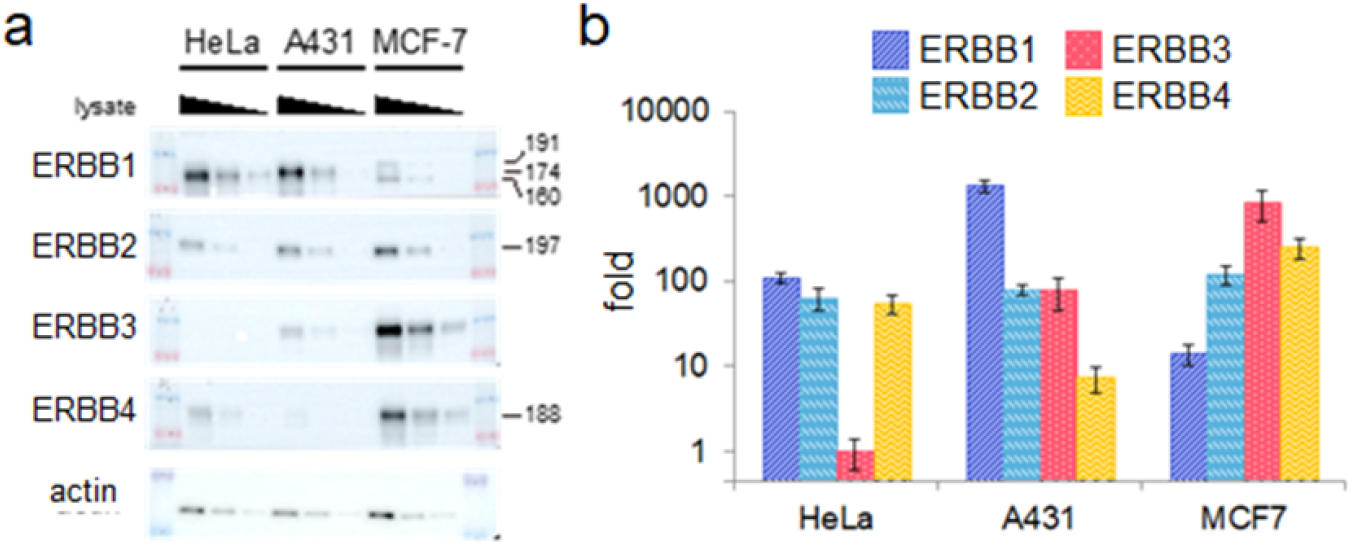
ERBB expression profiles in the three cell types. **a**. Western blotting of ERBB proteins. The lysate of each cell line was diluted in three steps to confirm the linearity of the antibody staining. **b**. Quantification of the ERBB expression level. Values were normalized to the ERBB3 expression in HeLa cells.

The time-course results indicated that the phosphorylation level changed dramatically within 5 min and then slowly changed the next 55 min. To focus on the initial responses directly induced by the ERBB–ligand interaction, we analyzed ERBB phosphorylation at 5 min, at which point feedback effects from downstream signaling are minimal (17).

The three cell types showed distinct behaviors in their phosphorylation dynamics to the same ligand, and the two ligands induced distinct dynamics in the same cell type.

### ERBB expression profile

The relative expression levels of ERBB1-B4 in HeLa, A431, and MCF7 cells were measured by western blotting (Fig. 3a). The densities of the observed bands for anti-ERBB antibodies were compensated for differences in the antibody titers (Fig. S1) to compare the expression levels (see Methods). The estimated ERBB expression levels were normalized to the lowest expression level of ERBB (HeLa ERBB3) (Fig. 3b).

The observed tendency of the expression profiles was similar to previous studies: ERBB1 was abundantly expressed in A431 cells (18), and ERBB3/ERBB4 were expressed more than ERBB1 in MCF7 cells (19).

### ERBB overexpression profile

To examine the validity of the estimated reaction rate constants, we tried to predict cell behavior after manipulating the ERBB expression level. We examined the following four conditions: (1) ERBB2 overexpression in HeLa cells (HeLa+B2), (2) ERBB3 overexpression in HeLa (HeLa+B3), (3) ERBB3 overexpression in A431 cells (A431+B3), and (4) ERBB1 overexpression in MCF7 cells (MCF7+B1). These overexpressions were chosen based on our hypothesis that increasing the expression levels of initially lowly expressed genes may induce large changes in the phosphorylation dynamics.

The overexpression levels of ERBB were measured by western blotting and found to be 51-, 260-, 61-, and 46-fold higher for HeLa+B2, HeLa+B3, A431+B3, and MCF+B1, respectively, compared with the expression levels in parental cells (Fig. 4a, upper). We confirmed that our measurements were not mixtures of cells by immunofluorescence, which showed that almost all cells (> 90%) overexpressed the transfected ERBB (Fig. 4a, lower). The phosphorylation of ERBBs in wild-type and ERBB-overexpressing cells was quantified using western blotting (Fig. 4b). Based on the overexpression levels quantified experimentally, we calculated the phosphorylation dynamics of ERBB, as described in the following section, and compared the phosphorylation levels between mutants experimentally and mathematically (Fig. 4c).

**Fig. 4.**
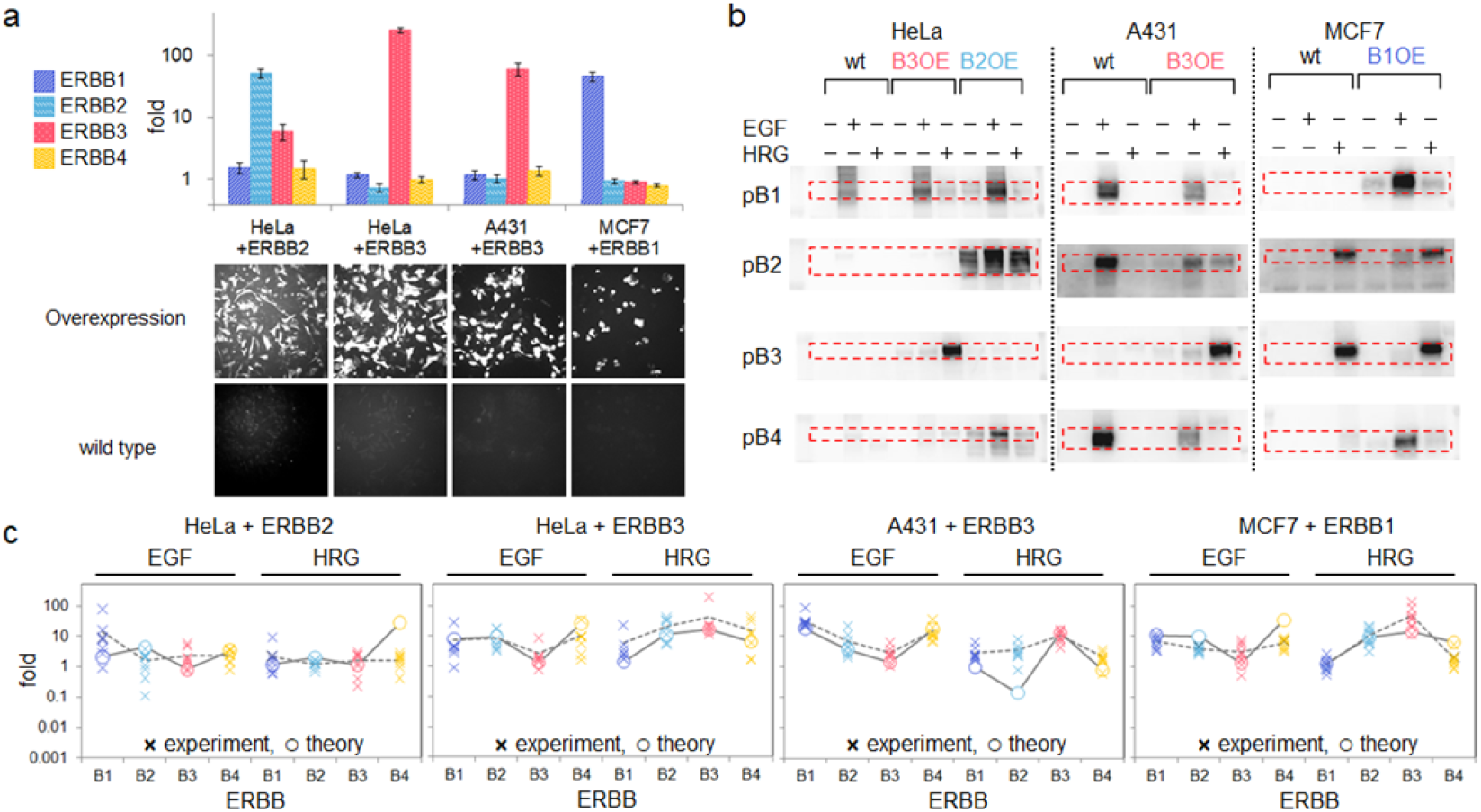
ERBB phosphorylation in cells overexpressing ERBB. **a**. ERBB expression levels (upper) and immunofluorescence images of overexpressed ERBB (lower) in the indicated cells. The expression level of each ERBB was normalized to that in wild-type cells. **b**. EGF or HRG induced ERBB-phosphorylation in wild-type (wt) and ERBB-overexpressing (OE) cells. The red dotted rectangles indicate phosphorylated ERBB bands. **c**. Fold change of the ERBB phosphorylation. Crosses and open circles respectively show experimental observations and theoretical predictions using parameters obtained from the global model fitting in Fig. 2c.

### Mathematical Analysis Mathematical modeling

In order to understand the dynamic changes of the phosphorylation levels, we constructed a model based on mass-action differential equations of chemical reactions (see (20–22) for studies based on mass action kinetics). We labeled the ERRB subtypes as *i* = 1,2,3,4. The 4 subtypes can be phosphorylated, ligand-bound (for *i* = 1,3,4; we do not consider ligand − bound ERBB2), or both, thus giving 14 monomer states in total, which we write as 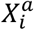 (*a* = bear, liganded, phosphorylated, both). These monomers form dimers 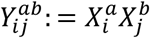 (*a, b* = bear, liganded, phosphorylated, both) (*i, j* = 1,2,3,4), such that there are 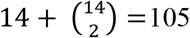 dimer states. In the early stage of the phosphorylation dynamics (≅ 5 min), there are three different classes of reactions: 1) ligand-binding and dissociation, 2) dimerization and dissociation, and 3) phosphorylation and dephosphorylation (Fig. 1a). We do not consider the endocytosis or degradation of ERBBs in this early stage of the signal transduction at 25 °C.

In order to reduce the number of variables and parameters in the model, we assumed that the ligand concentrations are negligible before the stimulation and that the ligand concentrations are sufficiently high and all ERBB proteins are ligand-bound after the stimulation. To validate this assumption, we experimentally confirmed that phosphorylation levels after stimulation are not sensitive to ligand concentrations, which indicates that the receptors were saturated with ligands in our experiments.

Thus, we can reduce the system size, since it is sufficient to consider either ligand-unbound states (before stimulation) or ligand-bound states (after stimulation), and we do not need to deal with their coexistence. More specifically, the concentrations were determined from the following equations:

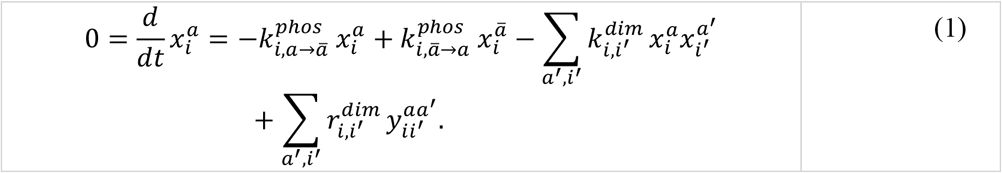

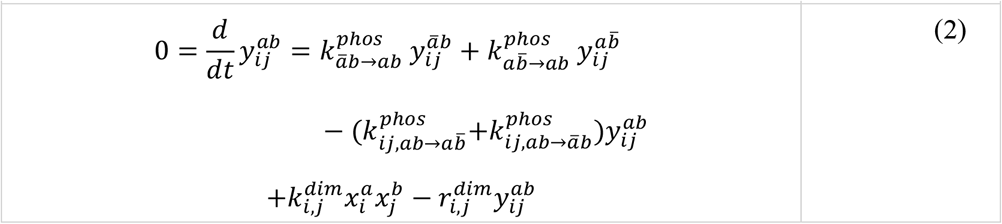

Here, 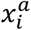 and 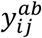 are the concentrations of the monomers 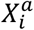 and dimers 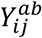, respectively. The indices *a, b* take only two states, {0=unphosphorylated, 1=phosphorylated}, and 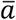 represents the other state, i.e., 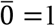 and 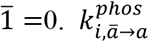 and 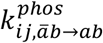 denote the phosphorylation/dephosphorylation rates of the monomers and dimers, respectively. For example, 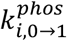 denotes the phosphorylation rate of ERBB*i* monomer, while 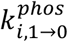 denotes the dephosphorylation rate. We assume that ERBB3 receptors are incapable of phosphorylating themselves and other receptors, namely 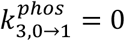 and 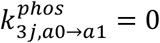. Finally, 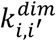 and 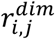 denote the dimerization rates and dissociation rates, respectively. An important point of our analysis is that we consider the steady-state equations (1) and (2) both before and after ligand stimulation, and the effects of the stimulation are implemented by using different values of the dimerization/dissociation rate constants and the dimer phosphorylation rate constants before and after the stimulation.

The dynamics have four conserved quantities: the total amounts of ERBB1, ERBB2, ERBB3 and ERBB4. We measured these four quantities experimentally for each cell type. Given these conserved quantities and the reaction rate constants, Eq. 1 and Eq. 2 determine the steady-state concentrations of all chemicals uniquely. In order to obtain a minimal model, we assumed that all cell types have the same reaction rate constants. As we show below, the diversity of the responses to a ligand stimulation among the cell types can be explained from the difference in the expression profiles of the ERBB receptors.

### Parameter estimation (3-cell fitting)

In order to estimate the parameter values of the reaction rate constants, we performed a Markov Chain Monte Carlo simulation (see Supplementary Materials for the implementation). For a given set of parameter values for each cell type, we computed the phosphorylation levels of ERBB1, ERBB2, ERBB3, and ERBB4 under three external environments, namely in the absence of ligands, in the presence of EGF and in the presence of HRG. Then we compared the obtained phosphorylation levels with those measured experimentally. Through the Monte Carlo simulation, we constructed an ensemble of parameter sets that explains the experimental results with some accuracy. For an “energy function” in the Monte Carlo simulation, we used the sum of squared errors of the phosphorylation levels,

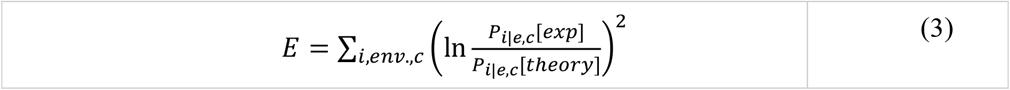

where the index *i*=1,2,3,4 labels the ERBB subtype, *e* labels the three environmental conditions, and *c* labels the three culture cell types (HeLa, MCF7, A431). *P*_*i*|*e,c*_[*exp*] and *P*_*i*|*e,c*_[*theory*] denote the experimentally and theoretically measured phosphorylation level of the ERBB *i* receptor of cell *c* under environment *e*. Note that *P*_*i*|*e,c*_ is the total concentration of phosphorylated ERBB *i* molecules, therefore, both phosphorylated monomers and dimers contribute to *P*_*i*|*e,c*_.

Fig. 2C shows the fold change in phosphorylation levels of ERBB receptors for two different ligands and three cell types. We estimated the parameter values of the model by the Monte Carlo method. The mathematical model explains the diversity of the phosphorylation levels of the ERBB receptors very well based on the difference in the receptor expression profiles.

### Validity the 3-cell fitting model

We calculated the phosphorylation dynamics of ERBB under the experimentally determined values of the total amounts of ERBB. The reaction rate constants were assumed to be the same for all cell types including the three wild-types and four overexpression mutants. The steady states of the ERBB phosphorylation obtained from the mathematical model are shown in Fig. 4c. Fig. 4c also shows experimental measurements of the phosphorylation levels at five minutes after the ligand stimulation.

The phosphorylation levels predicted by the mathematical model are close to the experimental measurements except for ERBB4 in HeLa+B2 mutants and ERBB2 in A431+B3 mutants in response to the HRG signal. In wild-type HeLa cells, large differences in the responses were observed among different ERBB subtypes to both EGF and HRG, i.e. ERBB4 phosphorylation was higher than ERBB3 phosphorylation. The mathematical model predicted the same tendency between ERBB4 and the other three subtypes to HRG in HeLa+B2 cells, although the responses of the four subtypes were almost the same experimentally. For the response to HRG in A431+B3 cells, the mathematical model shows that the ERBB2 response was lower than the ERBB3 response, however, the experimental difference of the response was much less.

### Re-estimation of reaction parameters (7-cell fitting)

To obtain higher validity, we re-estimated the reaction rate constants by integrating the observed phosphorylation levels of HeLa+B2, HeLa+B3, A431+B3, and MCF7+B1 in addition to HeLa, A431 and MCF7 cells into the fitting. A comparison of the experimental and new fitting results is shown in Fig. 5. The phosphorylation profiles obtained from the mathematical model were consistent with the observed profiles of the overexpression mutants as well as the wild-type cells. In the new fitting, we do not see the tendency of the deviation. We expect the estimated reaction rate constants to reflect the real phosphorylation dynamics more accurately than the 3-cell fit.

**Fig. 5.**
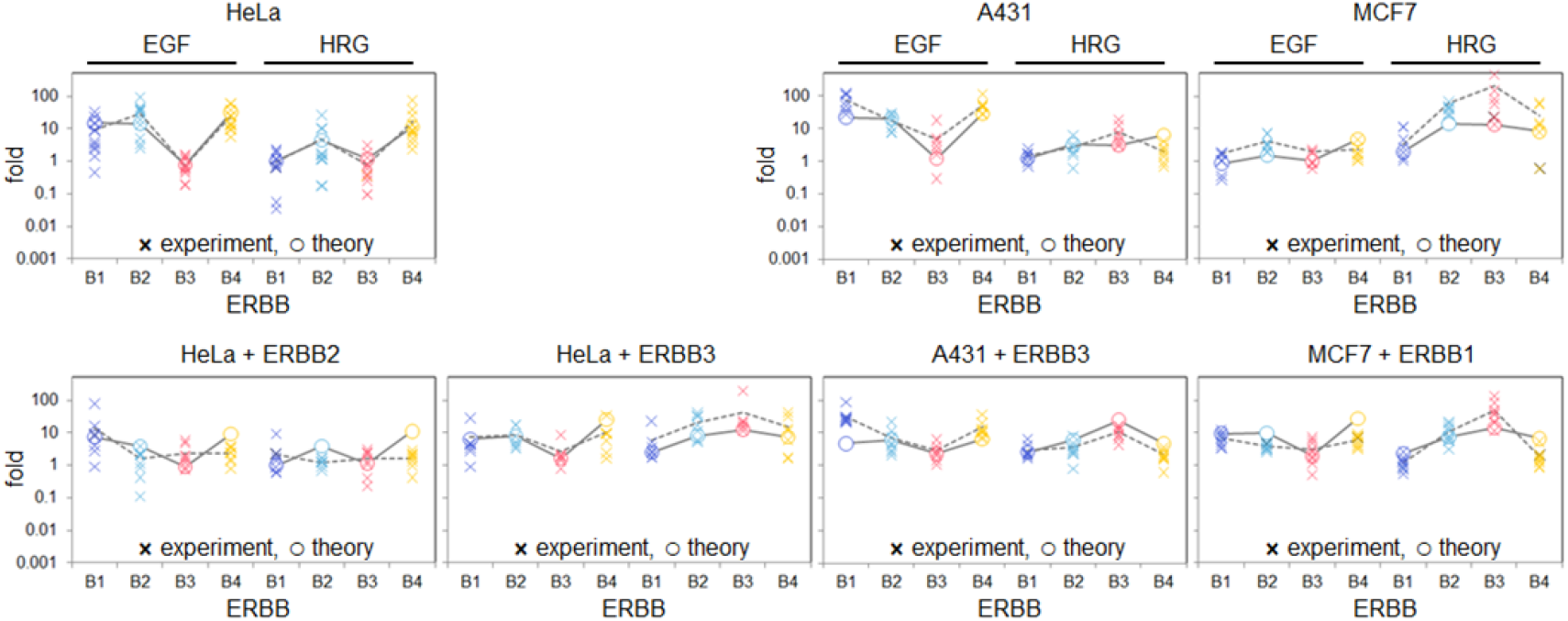
Experimental and mathematical phosphorylation levels. ERBB phosphorylation levels in ERRB wild-type (upper three panels) and overexpressing mutant (lower four panels) cells. Crosses and open circles respectively represent the experimental results and global model fitting. Bold crosses indicate experimental averages connected with dashed lines.

In the following, we discuss the characteristics of the estimated reaction rate constants in view of the biological network response. The average estimated values of the reaction rate constants are shown in table S1 and graphically represented in Fig. 6. The figure shows that the phosphorylation and dimerization rates depend on the combination of subtypes in the pairs. Affinities between the EBBB protomers are strong, especially in B1 and B4 homodimers even in the absence of any ligand. This prediction is consistent with the experimental results, indicating the presence of ERBB predimers (9, 10). The basal phosphorylation levels, which are determined by the balance between spontaneous phosphorylation and dephosphorylation reactions, are low in homodimers. As a result, basal phosphorylation in the ERBB system is maintained at a low level despite the presence of predimers. Dephosphorylation, but not phosphorylation, primarily determines the basal phosphorylation levels, which is consistent with previous reports (23, 24). The effect of ligand binding is highly variable among ERBB subtypes, e.g., EGF stimulates kinase activity but has little effect on the substrate activity of ERBB1, whereas HRG increases the substrate activity of ERBB3 and B4 as well as the kinase activity of ERBB4. This prediction must be verified with future experiments, however.

**Fig. 6.**
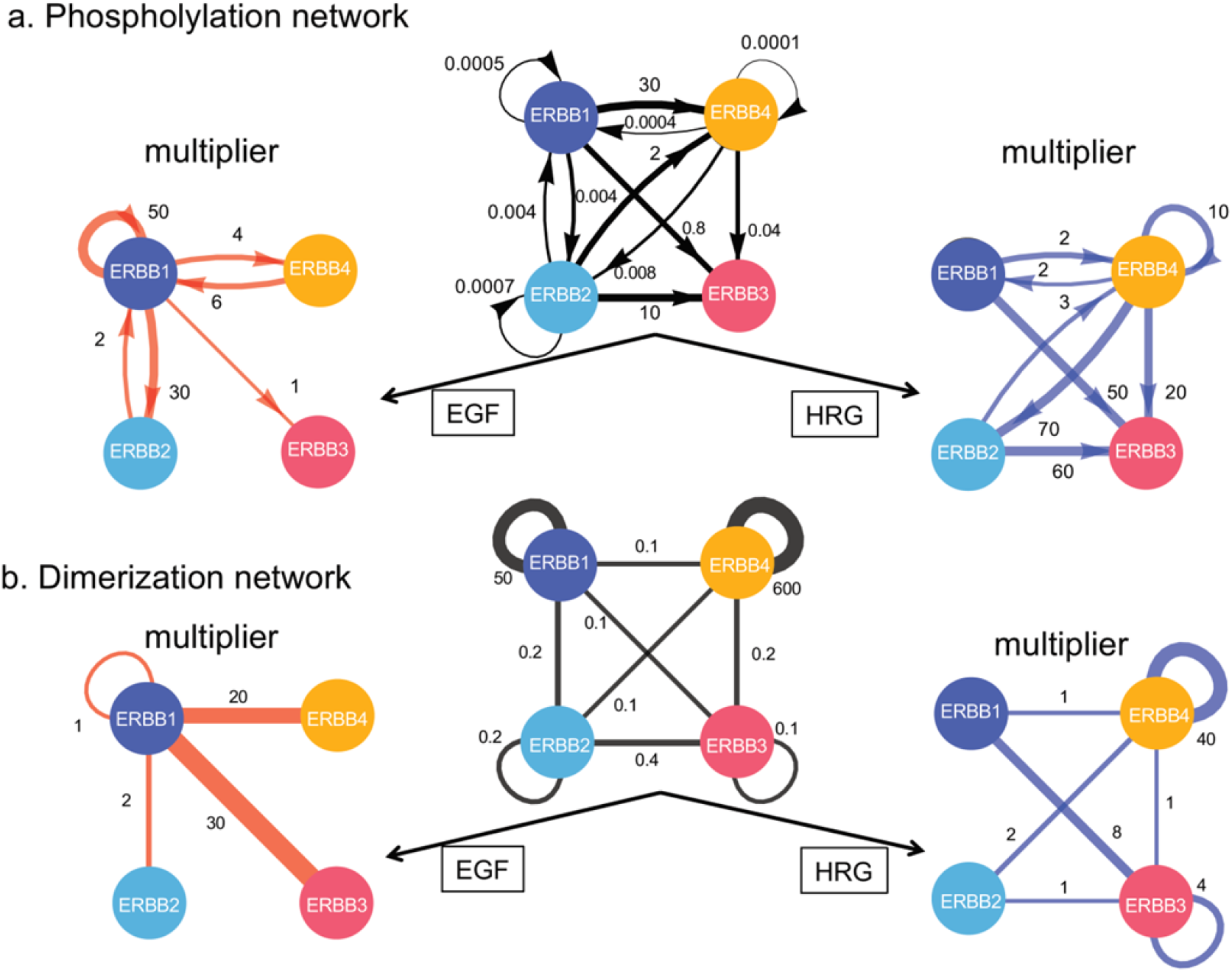
Ligand effects on the phosphorylation and dimerization networks. **a**. Phosphorylation networks. The black network in the middle shows effective phosphorylation rates on the edges, each of which is defined as the ratio of the dimer phosphorylation rate to the dephosphorylation rate. The left (red) and right (blue) networks represent multipliers on the effective rates due to EGF and HRG stimulation, respectively. **b**. Dimerization networks. The black network in the middle shows effective dimerization rates on the edges, each of which is defined as the ratio of the dimerization rate to the dephosphorylation rate. The left (red) and right (blue) networks represent multipliers on the effective rates due to EGF and HRG stimulation, respectively.

### Origin of the diversity of the phosphorylation dynamics

As we showed above, the ERBB phosphorylation dynamics strongly depend on the ERBB expression profiles. However, different ERBB expression profiles cannot induce large differences in dynamical behaviors if the four subtypes have similar kinetic properties (binding, dimerization and phosphorylation rates). Therefore, the diversity of the dynamical behaviors should depend both on the expression profiles and reaction rates of ERBBs. In the following, we study the diversity of the phosphorylation dynamics by distinguishing the diversity originated from the expression profiles, which we call “dose-dependent diversity”, and from the reaction rates, which we call “rate-dependent diversity”.

### Dose-dependent diversity

We examined the diversity of the phosphorylation dynamics by changing the ERBB expression patterns in the mathematical model. We prepared many expression profiles by setting three different expression levels (20, 100, 500) for each of the four subtypes, where all expression levels were normalized to ERBB3 in HeLa cells. (Discretizing the space of the expression levels into more than three levels does not change the results qualitatively.) For each of 81(= 3^4^) hypothetical expression patterns (shown in the left-end column in Fig. 7a), we simulated the phosphorylation dynamics and determined the phosphorylation levels before and after the ligand stimulation. The heat-map labeled “Unsym.” in Fig. 7a shows the fold change in phosphorylation levels of ERBB1-4 induced by EGF and HRG (columns) for each of the 81 ERBB expression patterns (rows). The pattern in the heat-map visualizes the diversity of the responses due to differences in the expression profiles, namely, dose-dependent diversity.

**Fig. 7.**
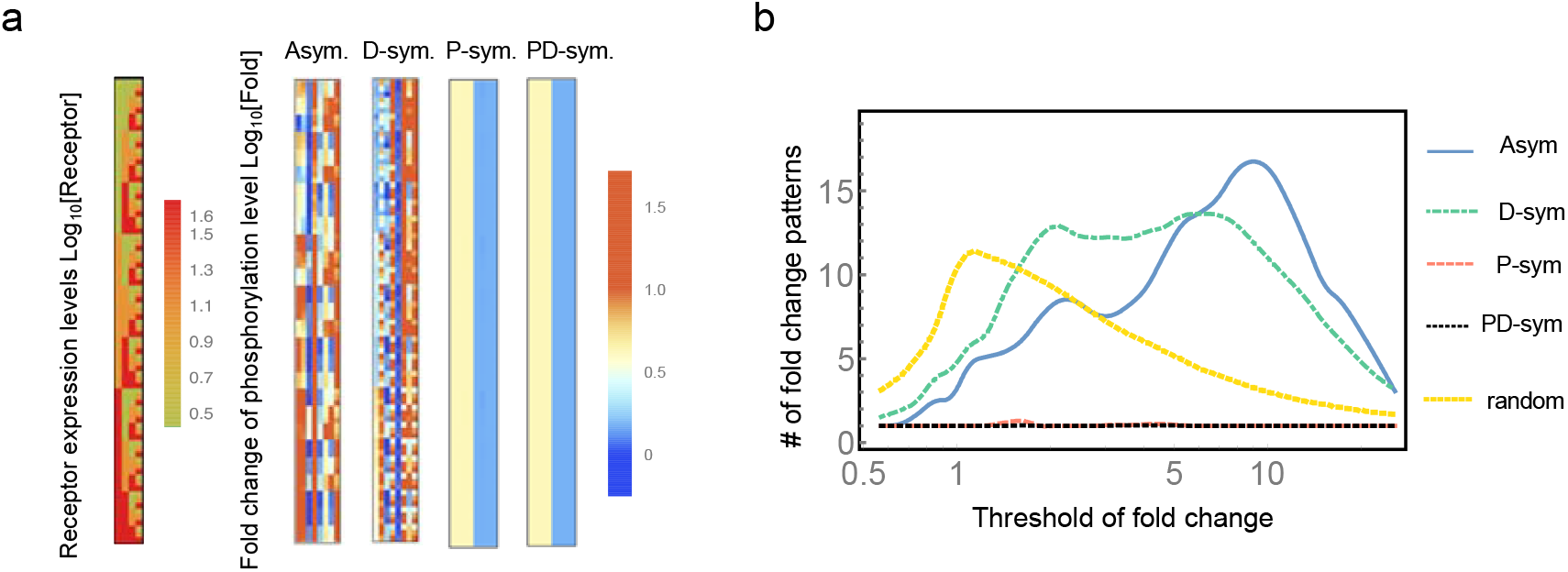
The diversity of phosphorylation patterns. a: Phosphorylation patterns obtained from the originally estimated parameters (unsymmetric kinetics; Unsym), symmetrized dimerization-associated parameters (D-sym), symmetrized phosphorylation-associated parameters (P-sym), and symmetrized phosphorylation- and dimerization-associated parameters (PD-sym). b: The response number against the threshold value used to discretize the fold change for Unsym, D-sym, P-sym, PD-sym and random kinetics.

In the heat-map in Fig. 7a, the responses of the four subtypes to two ligands (EGF and HRG) under 81 expression profiles are shown by continuous values. Each element of the 81×8 table is binarized depending on whether the element is higher or lower than a given threshold. Then, each row (the response for each ERBB expression profile) is represented by an 8-digit binary number, e.g. 01011011. Second, we count the number of different 8-digit numbers within the 81 ERBB expression profiles. We call these numbers “response numbers” to quantify the diversity of the phosphorylation patterns caused by differences in the ERBB expressions.

The response number depends on the threshold value used for the binarization. The curve labeled “Unsym.” shows the response number against the threshold value (Fig. 7b). The threshold of the phosphorylation level can be interpreted as the inability to detect phosphorylation (insensitivity) by the downstream molecules in the signal pathway. To characterize the curve, we also calculated the response number by using randomly sampled parameter values (Fig. 7b). More specifically, we sampled random values for [0.00001,1] reaction rates and [1,100] multipliers of the ligand stimulations.

The response number curve for the randomized model had a peak around the threshold with a fold change equal to 1. Note that under a setting of a threshold close to 1 (high sensitivity), phosphorylation responses to the ligands and spontaneous fluctuations cannot be distinguished by the downstream pathway. On the other hand, the response number curve obtained from the estimated parameter values (Fig. 7b) has a single peak at the threshold of about 10. This result implies that the phosphorylation dynamics of ERBB in human cells realizes the highest diversity of responses at a threshold of fold change 10, which seems to be sufficiently high to distinguish responses to ligands and spontaneous fluctuations.

The maximum value of the response number was less than 20, which is small compared to the 8-digit binary number (256=2^8^). Note that even if we take a wider domain of the expression level space and discretize the domain using a larger number of points, the response number does not change drastically (Fig. S2). This result together with Fig. 7b suggests that the upper bound of the diversity of the phosphorylation responses comes from the mathematical framework of the phosphorylation reaction, Eq. 1 and Eq. 2, rather than the choice of parameters. We expect that the number of different expression profiles of ERBB in human cells is also not so large, because large differences in expression profiles do not always drive large diversity in dynamical behaviors.

### Rate-dependent diversity

In this section, we discuss how differences in reaction rates (dimerization and phosphorylation) between different ERBB subtypes contribute to the diversity of the phosphorylation dynamics.

We first constructed “symmetric models”, where some reaction rate constants and the effects of the ligand stimulation were set to be equal among the different subtypes. More specifically, we considered three differently symmetrized models, D-symmetric, P-symmetric and PD-symmetric, wherein particular subsets of reaction rate constants were set to be equal among the different subtypes (Table S1). In the D-symmetric model, the values of the reaction parameters related to dimerization (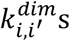 and 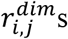, and multipliers of the ligand stimulation, respectively) were set to be geometric averages of the four subtypes in the original Un-symmetric model. Note that the symmetry of the multipliers for ligand binding implies uniformization of the ligand affinities between ERBBs. Similarly, in the P-symmetric model, the reaction rate constants related to phosphorylation (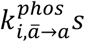 and 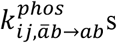) were set to be geometric averages of the original unsymmetric model. In the PD-symmetric model, four sets of parameters related to dimerization and phosphorylation (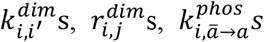 and 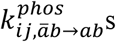) were set to be geometric averages. For each symmetric model, we calculated the phosphorylation dynamics and determined the steady states for all 81 expression profiles as done for the dose-dependent diversity analysis. We also calculated the response numbers for each symmetric model. From the decreases in the diversity in the symmetric models compared to the original unsymmetric model, we discuss the effects of the difference of the reaction rates to the diversity of the phosphorylation dynamics.

The heat-maps labeled “D-sym”, “P-sym” and “PD-sym” in Fig. 7a represent the diversities of the phosphorylation responses in each symmetric model. In the P-symmetric and PD-symmetric models, the color patterns in the heat-maps are much simpler than that in the unsymmetric and D-symmetric models.

In Fig. 7b, the response numbers against the threshold in the symmetric models are plotted on the same chart with the unsymmetric model. The smaller response numbers in the symmetric models compared to the unsymmetric model shows the effect of the variation in the reaction rate constants on the diversity of the phosphorylation responses (rate-dependent diversity). The response number of the D-symmetric model is comparable with the unsymmetric model. On the other hand, in the P-symmetric and PD-symmetric models, the response numbers are significantly lower. This result implies that the diversity of the phosphorylation rate constants between the four ERBB subtypes makes a much larger contribution to the diversity of the phosphorylation dynamics than the other dimerization parameters.

## Discussion

Here we studied the phosphorylation dynamics of ERBB receptors, which initiate multiple signal transduction pathways, by observing mutant cell lines experimentally and mathematically. We characterized the dynamics of the four ERBB subtypes as homodimers and heterodimers with different binding and phosphorylation rates. From our analysis, we found a diversity of phosphorylation responses depending on both the expression profiles and kinetic properties of the subtypes. Differences in the expression profiles may be a possible way to induce diverse responses among organs in the human body. Consistently, different expression profiles have been reported in different organs in the human body (Fig. 1c) (14–16).

Using our mathematical analysis, we examined the origin of the diversity of the phosphorylation responses of ERBBs to intercellular signals by decomposing the responses into dose-dependent diversity and rate-dependent diversity responses. We first confirmed that differences in the ERBB expression profiles induce the diversity of the phosphorylation responses. We found that under the parameter condition estimated from human cell lines, the maximum diversity of responses was realized when the sensitivity of the downstream molecules was sufficiently low (high in the threshold of the phosphorylation level) for distinguishing spontaneous fluctuations from responses to ligands. On the other hand, we found that the diversity of the phosphorylation responses cannot be larger than an upper bound based on the reaction dynamics even if the difference in the expression profiles is very large. Thus, these results suggest that the number of ERBB expression profiles in human cells is not so large, because a larger difference does not imply more diversity in dynamical behaviors.

Second, we showed that different reaction rates between subtypes is necessary for the diversity of the phosphorylation responses. From the analysis, the difference in phosphorylation rate constants between ERBB subtypes has a much larger contribution to the diversity of the phosphorylation responses than do the rate constants for dimerization. This finding implies that the divergent evolution in phosphorylation reactions after the gene duplications was essential to generating the diversity of phosphorylation responses among ERBB subtypes. The C-terminus phosphorylation domain, which is the substrate of the ERBB kinase domain, shows a much larger sequence difference than other domains. This difference may cause the difference in the phosphorylation rate constants among ERBB subtypes.

The decomposition of dose-dependent diversity and rate-dependent diversity responses in the model are difficult to study experimentally. For the dose-dependent diversity analysis, we calculated the number of patterns in the phosphorylation responses under exhaustive changes in the ERBB expression profiles. Experimentally, this type of analysis is not practical. As for the rate-dependent diversity analysis, we examined the phosphorylation responses under the symmetry of some reaction rate constants. This examination cancels the difference in the reaction properties between ERBB subtypes obtained by evolutionary divergence. These experiments too are not practical. By integrating experimental and theoretical methods, we obtained a deeper understanding of the behavior of the ERBB system from extensive and evolutionary analyses.

## Author Contributions

T.O. conceived and carried out all of the mathematical analyses. H.M. carried out the experiments. Y.S. directed experimental analysis and discussions. M.H. designed and carried out the experiments. A.M. organized the project. A.M., T.O., Y.S., and M.H. wrote the manuscript.

## Acknowledgement

We thank H. Sato and A. Kanayama for experimental support. This research was supported by the CREST program (grant no. JPMJCR13W6) of the Japan Science and Technology Agency (JST) (http://www.jst.go.jp/EN/index.html), the RIKEN iTHEMS Program, and the Joint Usage/Research Center Program of the Institute for Frontier Life and Medical Sciences, Kyoto University. YS was supported by MEXT Japan with Grants-in-Aid for Scientific Research (19H05647) and JST with CREST (JPMJCR1912). MH was supported by MEXT Japan with Grants-in-Aid for Scientific Research(B) (18H01839) and Grant-in-Aid for Scientific Research on Innovative Areas (18H05414). All Authors declare that there is no conflict of interest.

